# Preserved motor memory in Parkinson’s disease

**DOI:** 10.1101/2021.04.30.441882

**Authors:** Soraya Lahlou, Ella Gabitov, Lucy Owen, Daphna Shohamy, Madeleine Sharp

**Affiliations:** Department of Neurology and Neurosurgery, Montreal Neurological Institute, McGill University; Department of Psychological and Brain Sciences, Dartmouth College; Department of Psychology, Columbia University; Zuckerman Mind Brain Behavior Institute and Kavli Institute for Brain Science, Columbia University

**Keywords:** Parkinson’s disease, memory, consolidation, dopamine, motor sequence learning, cognition, aging

## Abstract

Patients with Parkinson’s disease, who lose the dopaminergic projections to the striatum, are impaired in certain aspects of motor learning. Recent evidence suggests that, in addition to its role in motor performance, the striatum plays a key role in the memory of motor learning. Whether Parkinson’s patients have impaired motor memory and whether motor memory is modulated by dopamine at the time of initial learning is unknown. To address these questions, we measured memory of a learned motor sequence in Parkinson’s patients who were either On or Off their dopaminergic medications. We compared them to a group of older and younger controls. Contrary to our predictions, motor memory was not impaired in patients compared to older controls, and was not influenced by dopamine state at the time of initial learning. To probe post-learning consolidation processes, we also tested whether learning a new sequence shortly after learning the initial sequence would interfere with later memory. We found that, in contrast to younger adults, neither older adults nor patients were susceptible to this interference. These findings suggest that motor memory is preserved in Parkinson’s patients and raise the possibility that motor memory in patients is supported by compensatory non-dopamine sensitive mechanisms. Furthermore, given the similar performance characteristics observed in the patients and older adults and the absence of an effect of dopamine, these results raise the possibility that aging and Parkinson’s disease affect motor memory in similar ways.

## 1. INTRODUCTION

It is well established that the striatum and striatal dopamine play a fundamental role in motor learning (Doyon et al., 2009a; Graybiel and Grafton, 2015; Yin and Knowlton, 2006). More recent evidence also suggests that the striatum – a subcortical structure that is richly innervated by dopaminergic neurons – plays a specific role in the memory consolidation phase of human motor learning (Debas et al., 2014; Doyon et al., 2003; Perrin and Venance, 2019; Pisani et al., 2005), but surprisingly little work has been done to establish the exact role that dopamine plays in this process. Parkinson’s patients, who lose the dopaminergic projections to the striatum, are impaired at certain aspects of motor learning. Recent evidence hints that they have an additional impairment in motor memory consolidation, but whether this relates directly to their dopaminergic loss is unknown (Dan et al., 2015; Doyon, 2008; Fernandes et al., 2017; Olson et al., 2019; Terpening et al., 2013). Meanwhile, if, and how, dopamine supports motor memory is of considerable clinical interest. Most patients with Parkinson’s disease, despite best efforts at pharmacologic dopamine replacement, spend considerable amounts of time in the low (‘off’) dopamine state. Establishing whether dopamine supports long-term memory for motor skills would determine whether the consequences to a patient of being in a low or high dopamine state actually extend beyond the typically considered short window of medication effect.

Consolidation is generally defined as an active and time-dependent process during which initially labile memories are strengthened and rendered resistant to interference (Dudai et al., 2015; Walker et al., 2003). In the case of motor memory consolidation, a behavioural hallmark of this process is the offline improvement of performance (Fischer et al., 2002; Robertson et al., 2004). Multiple lines of evidence using animal models suggest that dopamine plays a role in memory consolidation. For example, in rodents, pharmacologic studies have shown that, across different forms of memory, manipulating dopamine around the time of the initial learning of a task influences memory consolidation for that task (Bethus et al., 2010; McNamara et al., 2014; Takeuchi et al., 2016; White et al., 1993). In humans, functional neuroimaging studies have provided indirect support for a role of striatal dopamine in motor consolidation by showing that changes in striatal BOLD activity during the initial learning are predictive of the off-line improvement in performance (Albouy et al., 2015, 2013a; Debas et al., 2010; Pinsard et al., 2019). It has also been shown that humans retain a motor skill better if they are provided with reward at the time of the initial learning of that skill –a behavioural manipulation thought to indirectly engage the dopaminergic system (Abe et al., 2011).

These findings raise the possibility that striatal dopamine loss in Parkinson’s patients may lead to impaired motor memory. Aging may also impact memory in Parkinson’s patients. Indeed, older adults do not show the offline improvements typically seen in younger adults, though they do maintain their performance across a delay, indicating that some aspects of a motor memory are nonetheless consolidated (Spencer et al., 2007; Wilson et al., 2012). Though most studies on motor learning in Parkinson’s patients have focused on the initial acquisition of a motor skill rather than subsequent memory consolidation of that skill (Clark et al., 2014; Hayes et al., 2015; Ruitenberg et al., 2015), two recent studies have shown an absence of offline gains in Parkinson’s patients, thereby providing preliminary evidence for the presence of impaired motor memory consolidation (Dan et al., 2015; Terpening et al., 2013). However, neither of these studies manipulated dopamine medications. It therefore remains unknown whether the presence of dopamine at the time of initial acquisition of a motor skill plays a specific role in enhancing the subsequent memory of that skill. This has implications for understanding whether the motor memory deficits of Parkinson’s disease are distinct from those related to aging.

To address these questions, we measured motor memory consolidation in two groups of Parkinson’s patients, one tested On and one tested Off dopaminergic medications, and compared them to healthy older controls. We also validated our task design in a group of young controls. All participants were initially trained to repeatedly tap a 5-element sequence (Doyon, 2008; Doyon et al., 2003; Korman et al., 2007; Pinsard et al., 2019). Motor memory consolidation was assessed in two ways: 1) we measured the change in performance across a two-day delay, and 2) in separate groups of participants, we also measured how these changes were affected by an interference manipulation. The interference manipulation consisted of having participants learn a second, different sequence two hours after the initial acquisition (Korman et al., 2007), which allowed us to probe whether susceptibility to interference is influenced by dopamine state. We hypothesized that dopamine at the time of initial acquisition would positively affect the consolidation phase of learning and lead to better motor memory at the delayed retest.

Specifically, we predicted that patients On dopaminergic medications would exhibit better maintenance of performance across the delay and reduced susceptibility to interference compared to patients Off medication.

Contrary to our predictions, we found that dopamine state at the time of initial acquisition did not influence motor memory maintenance. Overall, memory maintenance in the patients was similar to that of older adults and neither group showed offline gains. Interestingly, we also found that, unlike the young adults, neither older adults nor patients were susceptible to interference conducted two hours after initial learning. These results raise the possibility that motor memory in Parkinson’s patients relies on compensatory extra-striatal and non-dopamine-dependent mechanisms, and that these mechanisms may be similar to those that explain age-related differences in motor memory consolidation.

## 2. METHODS

### 2.1. Participants

Parkinson’s patients were recruited either from the Center for Parkinson’s Disease and other Movement Disorders at the Columbia University Medical Center or from the Michael J Fox Foundation Trial Finder website. Fifty-two patients were tested but 4 were excluded: of those, 2 patients did not meet the inclusion criteria for the task, which are detailed below, and 2 patients were excluded due to missing data. The analyses were conducted on 25 patients who were tested OFF their dopaminergic medications (PD-OFF; mean + SD age: 62.0 + 7.60, disease duration 6.6 + 3.9 years), and 23 patients who were tested ON their dopaminergic medications (PD-ON; mean + SD age: 63.2 + 7.0, disease duration: 6.8 + 3.24 years). All patients were receiving levodopa and endorsed levodopa responsiveness. In addition, 9/25 PD-OFF and 10/23 PD-ON were also being treated with a dopamine agonist. Participants had to use their non-dominant hand for the motor task. In the case of the PD-OFF, the non-dominant hand was also the less affected hand in 11/25. In the PD-ON it was the less affected in 11/23. Twenty-three healthy older controls were also tested (HC; Male 10, mean age 62.2 (SD = 7.52)). Demographic and clinical details are provided in **Table 1**.

**Table 1.** Demographic and clinical characteristics of participants retained for analyses. Table shows mean (SD). *P*-value from ^a^ANOVA for group differences or ^b^Pearson’s Chi-squared test. ^c^ MoCA = Montreal Cognitive Assessment; ^c^ Digit Span total = sum of forward and backward span; ^d^ UPDRS = Unified Parkinson’s Disease Rating Scale-Part III, tested ON in ON group, and OFF in OFF group; ^f^ LEED = Levodopa equivalent dosing, includes levodopa, dopamine agonists, amantadine, monoamine oxidase inhibitors and catechol-O-methyl transferase inhibitors

|  | Healthy controls<br>(n=23) | Parkinson's OFF<br>(n=25) | Parkinson's ON<br>(n=23) | *p*-value |
| --- | --- | --- | --- | --- |
| **Age** | 62.2 (7.52) | 62.0 (7.60) | 63.2 (7.02) | 0.837<sup>a</sup> |
| **Sex (male)** | 10/23 | 13/25 | 15/23 | 0.329<sup>b</sup> |
| **Education (years)** | 17.6 (2.40) | 18.9 (2.24) | 18.7 (2.76) | 0.204<sup>a</sup> |
| **MoCA <sup>c</sup>** | 28.2 (1.51) | 28.4 (1.59) | 28.3 (1.73) | 0.875<sup>a</sup> |
| **Digit Span total <sup>d</sup>** | 13.0 (2.07) | 12.3 (1.86) | 13.2 (1.91) | 0.272<sup>a</sup> |
| **UPDRS <sup>e</sup>** | n/a | 25.7 (8.76) | 23.2 (7.96) | 0.328<sup>a</sup> |
| **LEED (mg) <sup>f</sup>** | n/a | 630 (232) | 668 (318) | 0.636<sup>a</sup> |
| **Disease duration** | n/a | 6.60 (3.93) | 6.83 (3.24) | 0.829<sup>a</sup> |
| **Right-handed** | 19/23 | 18/25 | 20/23 | 0.395<sup>b</sup> |
| **Used less affected hand** | n/a | 10/25 | 11/23 | 0.798<sup>b</sup> |

To establish the validity of the task, thirty-four young controls were also tested (19 males, mean age 21.3 (SD = 4.37), 28 right-handed). None had active neurologic or psychiatric disease.

### 2.2. Medication manipulation

Parkinson’s disease patients were in the same drug state for both the acquisition phase of the task and the delayed memory test (i.e. ON dopaminergic medications for Day 1 and Day 3 or OFF for both; **Figure 1**). Details of the study design are provided below. Patients tested ON took their usual dose of medications 1 hour before the start of testing. Patients tested OFF dopaminergic medication were instructed to take their last dose the evening before the experiment (average time since last dose = 17 hours). We did not manipulate dopaminergic medications during the period between the two testing days, patients were told to take their medications as they usually do. In particular, we did not manipulate overnight dopaminergic intake (this applies only to Night 1; during Night 2, which preceded Day 3, patients were either ON or OFF according to their group assignment). As a result, some patients were receiving dopamine replacement during the period of sleep of Night 1, either from short-acting medications taken just before bed, or from longer-acting dopamine agonists taken in the afternoon or evening. This was the case for a similar proportion of patients in each group (11/25 PD-OFF and 12/23 PD-ON; p = 0.8). We conducted a supplementary analysis including only participants who did not receive any dopamine replacement overnight and showed that the pattern of results is similar to that of the full sample (Results).

**Figure 1.**
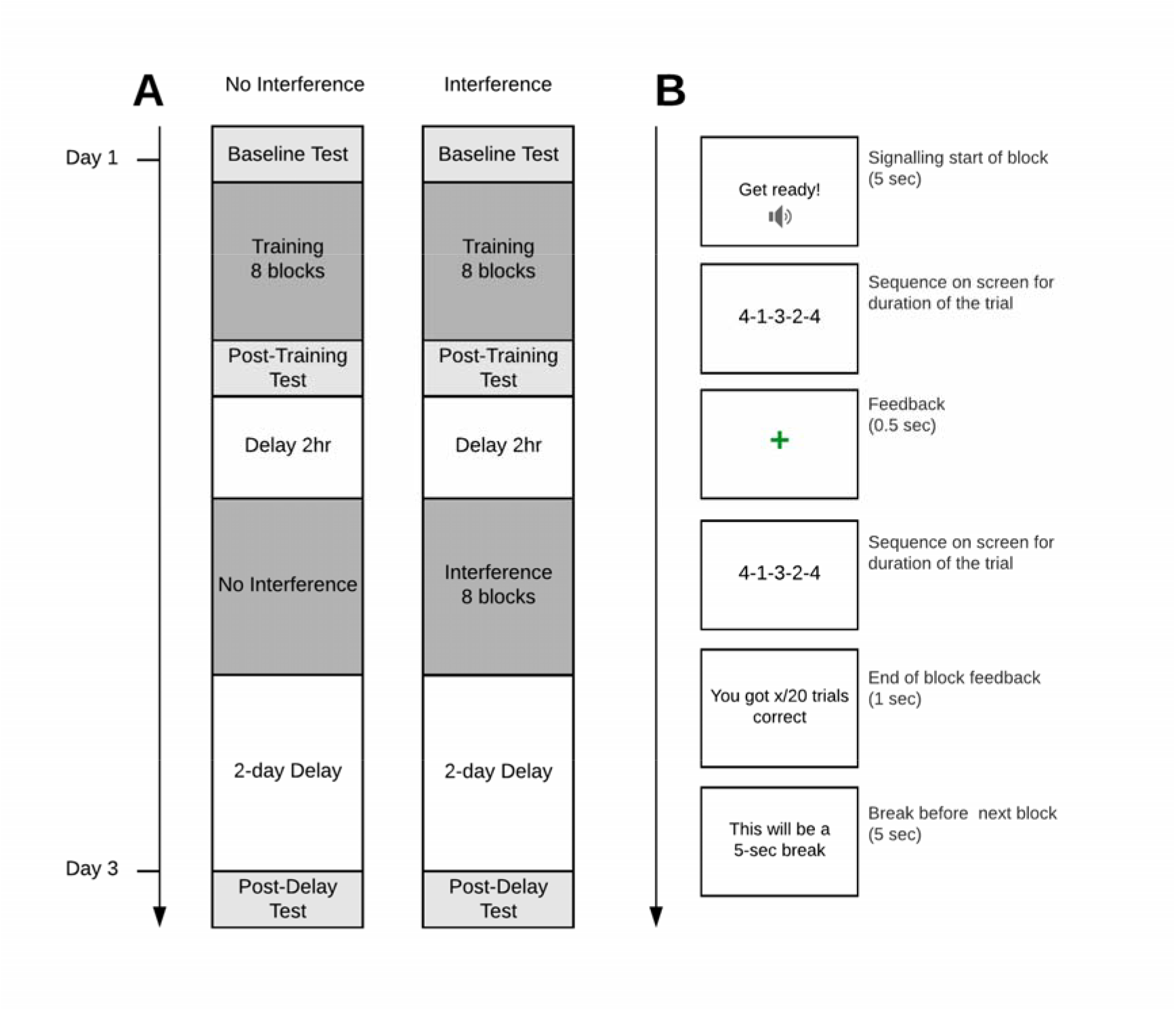
Schematic of the experimental design and trial sequence. (**A**) Participants were assigned to either the Interference or No interference condition. All groups were first assessed on their baseline performance (Baseline test). The test consisted of repeatedly tapping the target sequence ‘4-1-3-2-4’ as fast and accurately as possible for 4 trials of 30 seconds. Participants were then trained on the motor sequence over 8 blocks of 20 trials and received feedback on their performance. A Post-training test immediately followed. Two hours later, the Interference group was trained on a new sequence composed of the same elements but in reverse order (‘4-2-3-1-4’) for 8 blocks of 20 trials, whereas participants in the No Interference group did not undergo this procedure but remained in the lab for the same amount of time. All participants returned to the laboratory two days later and were tested on the original sequence (Post-delay test). Each of the three tests followed the same procedure. (**B**) Trial sequence during the training phase: a “Get ready!” screen accompanied by a sound signaled the beginning of a block, participants then performed the target sequence, which remained visible on the screen until 5 keys had been pressed. Feedback was given in the form of a green fixation cross for correct trials or a red fixation cross for incorrect trials. At the end of each block, the screen displayed how many sequences out of 20 were correctly performed, followed by a 5-second break prior to the next training block.

### 2.3. Motor sequence learning task

Participants performed a computerized version of the motor sequence learning task adapted from Karni et al. (1998, 1995). This task has been extensively used to study motor learning consolidation and its underlying processes in humans. Participants were trained to tap a 5-element sequence on a keyboard, as fast and accurately as possible, using 4 keys (F, G, H, J) which were labelled 1, 2, 3, 4 respectively. The sequence to learn was 4-1-3-2-4. All participants used their non-dominant hand and were instructed to use only 4-fingers, excluding the thumb, such that the pinky pressed the 1, the ring finger the 2, the middle finger the 3 and the index the 4 (in the case of left-hand use). The task consisted of a training phase and of three tests (baseline, post-training and post-delay; **Figure 1**). First, participants completed a baseline test of performance where they were given four 30-second trials during which they were required to repeatedly generate the entire sequence as many times as possible. Speed and accuracy were both emphasized. The numbers representing the sequence (i.e., “4-1-3-2-4”) remained on the screen at all times. Participants had a 30-second break between trials to rest their hand. Then, during the training phase, participants were given eight blocks of training during which they were required to perform the sequence 20 times. Participants received feedback for correct or incorrect performance after each series of 5 keys pressed in the form of a green or red fixation cross. A post-training test of performance, identical to the baseline test, immediately followed the training. Participants were randomly assigned to one of two conditions: the Interference condition or the No Interference condition. In the case of the Interference condition, two hours after the end of training, participants were trained to perform a new 5-element sequence composed of the same elements but in the reverse order (4-2-3-1-4), following the same procedure as the initial training. The two-hour delay was chosen because it has been shown that this interval is within a time-window during which the newly acquired motor skill is susceptible to interference (Korman et al., 2007). In the No Interference condition, participants stayed in the lab for the same amount of time but did not perform any particular task. During the 2-hour delay, both groups completed another task which has been reported separately (Sharp et al., 2020). All participants returned two days later for a post-delay test following the same procedure as the baseline test. The task was implemented using the PsychoPy2 Experiment Builder (v1.82.00) (Peirce, 2008, 2007).

### 2.4. Analysis

We excluded data from participants whose mean number of correct sequences per trial on the Post-training test was 2 SD away from their group means. We applied this criteria only to posttraining performance because it was used as a baseline to measure memory. As a result, one PD-ON and one PD-OFF patient were excluded completely. Another 1 HC scored zero on three out of the four trials at both the baseline and post-delay tests indicating poor task engagement and was thus also excluded. For the remaining participants, single trials from any of the test sessions with zero correct sequences were substituted with the average of the trial before and after, because it was assumed that these isolated instances reflected misuse of the keys rather than true poor performance. The removal of single trials occurred in 7 participants (2 HC, 3 PD-OFF and 2 PD-ON). If this occurred on more than one trial of the test session but affected only a single test, the whole test session was removed from the analysis; this occurred in 2 participants: 1 PD-OFF and 1 HC had a single test session removed. The other single instance when a test session was removed was when a participant significantly underused the allotted test time (one HC used only 2 of the 30 seconds allotted for a test trial on a baseline test). The final samples were as follows: 23 HC, 25 PD-OFF, 23 PD-ON and 34 Young controls.

In keeping with previous approaches to analyzing performance on this task, we computed a performance index which considered both accuracy (number of correct sequences performed) and duration (performance duration from the first key-press to the last key-press) as our main outcome measure used to establish levels of performance at each of the timepoints (Performance Index = number of correct sequences / duration * 10) (Dan et al., 2015). Computing a performance index allows us to account for the slight variations in the duration of a trial (although 30 seconds were allotted for test trials, few-second delays occasionally occurred prior to initiation of the first sequence). The performance index also accounts for errors since keys pressed in error necessarily reduce the time available to produce a correct key press. We also separately performed analyses using number of correct sequences and mean duration of correct sequences as outcome measures. These analyses are presented in the Supplementary materials.

To confirm that groups did not differ in their baseline performance nor in the magnitude of performance improvement derived from training, and groups assigned to the two interference conditions did not differ in their baseline performance, we ran a three-way mixed ANOVA on the performance index, with test as a within-subject factor, and group and interference condition as between-subject factors (Test [Baseline, Post-Training] x Group [HC, ON, OFF] x Interference [Int, NoInt]).

Our main analyses focused on examining the differences between groups in the maintenance of performance across the delay and in the susceptibility to interference. We ran a three-way mixed ANOVA on the performance index derived from the post-training test and the post-delay test, with test as a within-subject factor and group and interference condition as between-subject factors (Test [Post-Training, Post-Delay] x Group [HC, ON, OFF] x Interference [Int, NoInt]). We examined within group differences using two-sample t-tests. We also computed Bayes Factors for within group differences. Analyses were conducted in R version 4.0.5 using the ‘ez’ and ‘BayesFactor’ packages (Lawrence, 2016; Morey and Rouder, 2018).

## 3. RESULTS

### 3.1. Replication of previous findings in young adults

First, we validated we were able to replicate the findings typically observed in young adults. Specifically, it has been repeatedly shown in healthy young adults that 1) offline gains in performance occur over an overnight delay, and 2) that interference administered two hours after the initial learning leads to a reduction of offline gains (Doyon et al., 2009b; Korman et al., 2003; Robertson et al., 2004; Walker et al., 2002). As expected, the young healthy controls showed an offline improvement in performance after the two-day delay (mean of change in performance index: 0.83 units, 95% CI [0.61, 1.05], t= 8.05, df = 17, p < 0.005). Furthermore, we found a trend suggesting that participants who underwent interference after learning showed less offline-gains than those who did not (between-condition difference in offline-gains = −0.533, 95% CI [-0.02, 1.08], t= 2.00, df=20.24, p = 0.059) (**Supplemental Figure 1**).

### 3.2. Baseline performance and training gains in older adults and patients

Prior to the training, HC, PD-OFF and PD-ON successfully completed 12.9±0.77, 10.8±0.55, and 10.8±0.60 sequences during the baseline test, with a corresponding performance index of 4.47±0.25, 3.93±0.27 and 3.71±0.20, respectively (mean±s.e.m.). Baseline performance was comparable across groups (Group [HC, OFF, ON]: F(2, 62)= 0.94, p=0.39) and all three groups showed a similar improvement in performance following training (Group [HC, OFF, ON] x Test [Baseline, Post-Training]: F(2, 62)= 2.01, p=0.14; **Figure 2**). We found a similar pattern of performance when we examined number of correct sequences and sequence duration as outcome measures instead of performance index (Supplementary Figure 1 and 4).

**Figure 2.**
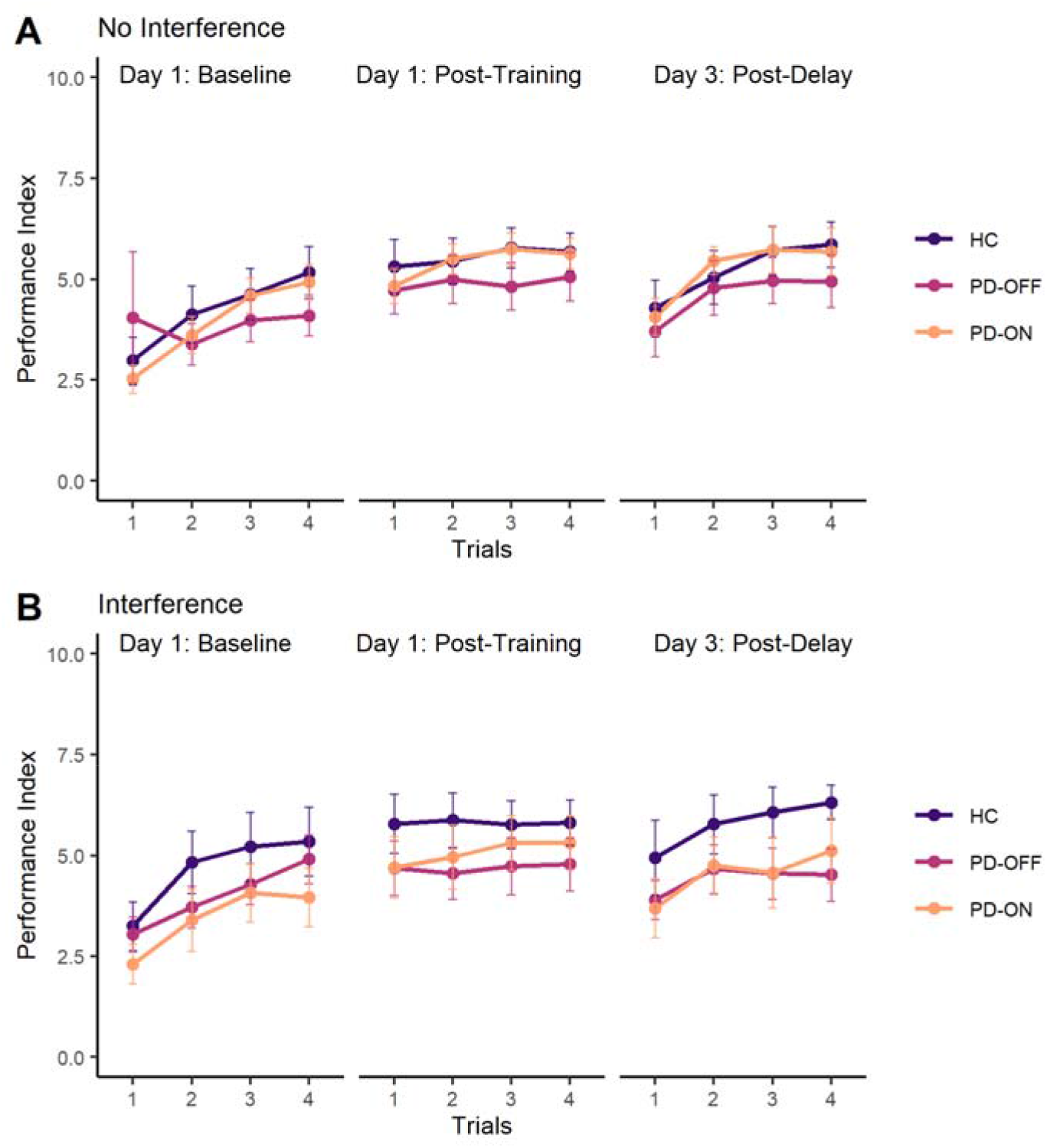
Test performance. Performance measured with the Performance index (i.e., number of correct sequences/duration * 10) is shown across trials (1-4) for each of the three test sessions (baseline, posttraining and post-delay) for all three groups (HC, PD-OFF and PD-ON). Conditions are presented separately: participants that did not undergo the interference (A) and those that did (B). Participants across groups and conditions show similar baseline performance, similar improvements from training and similar maintenance of performance across the two-day delay. Error bars represent s.e.m. within group and trial.

### 3.2. Motor memory consolidation in older adults and patients

We first hypothesized that the low dopamine state of Parkinson’s disease would affect motor memory, and that patients OFF would show worse maintenance of motor memory across the two-day delay than healthy controls and patients ON. However, using a repeated-measures ANOVA to compare post-training and post-delay performance, we found no group differences in the degree of maintenance of memory across the delay (Group [HC, OFF, ON] x Test [Post-Training, Post-Delay]: F(2, 65)= 0.32, p=0.72). In fact, unlike the young adults, none of the groups showed offline gains, and instead the was evidence for a slight decay in memory which was significant only in the controls and patients OFF (mean change in performance index: HC - 0.33 units, p=0.06; OFF −0.29 units, p=0.03; ON −0.19 units, p=0.32). We also hypothesized that patients OFF medication would show more susceptibility to interference than patients ON and healthy controls. However, we found no differences in the maintenance of performance when comparing participants who had undergone the interference to participants who hadn’t (Group [HC, OFF, ON] x Interference [NoInt, Int] x Test [Post-Training, Post-Delay]: F(2, 65)=1.00, p=0.37).

To examine the effects of interference more closely, we examined, within each group, the effect of Interference on the degree of performance maintenance across the delay. None of the groups showed a difference in motor memory maintenance related to the interference. Specifically, memory was not worse in participants who had undergone interference than in participants who had not (HC: t= −1.07, p=0.29; PD-OFF: t= −0.05, p=0.95; PD-ON: t=0.95, p=0.35) (**Figure 3**). Because the absence of an effect of interference on memory maintenance in all groups was unexpected, and because these within group comparisons were conducted by comparing smaller sub-samples, we additionally computed Bayes factors for each t-test to attempt to quantify the evidence for the null. In all three groups, the Bayes factor for the t-test examining the effect of interference on memory maintenance suggested weak support for the null hypothesis over the alternative (BF_HC_=1.70, BF_OFF_=2.71, BF_ON_=1.82).

**Figure 3.**
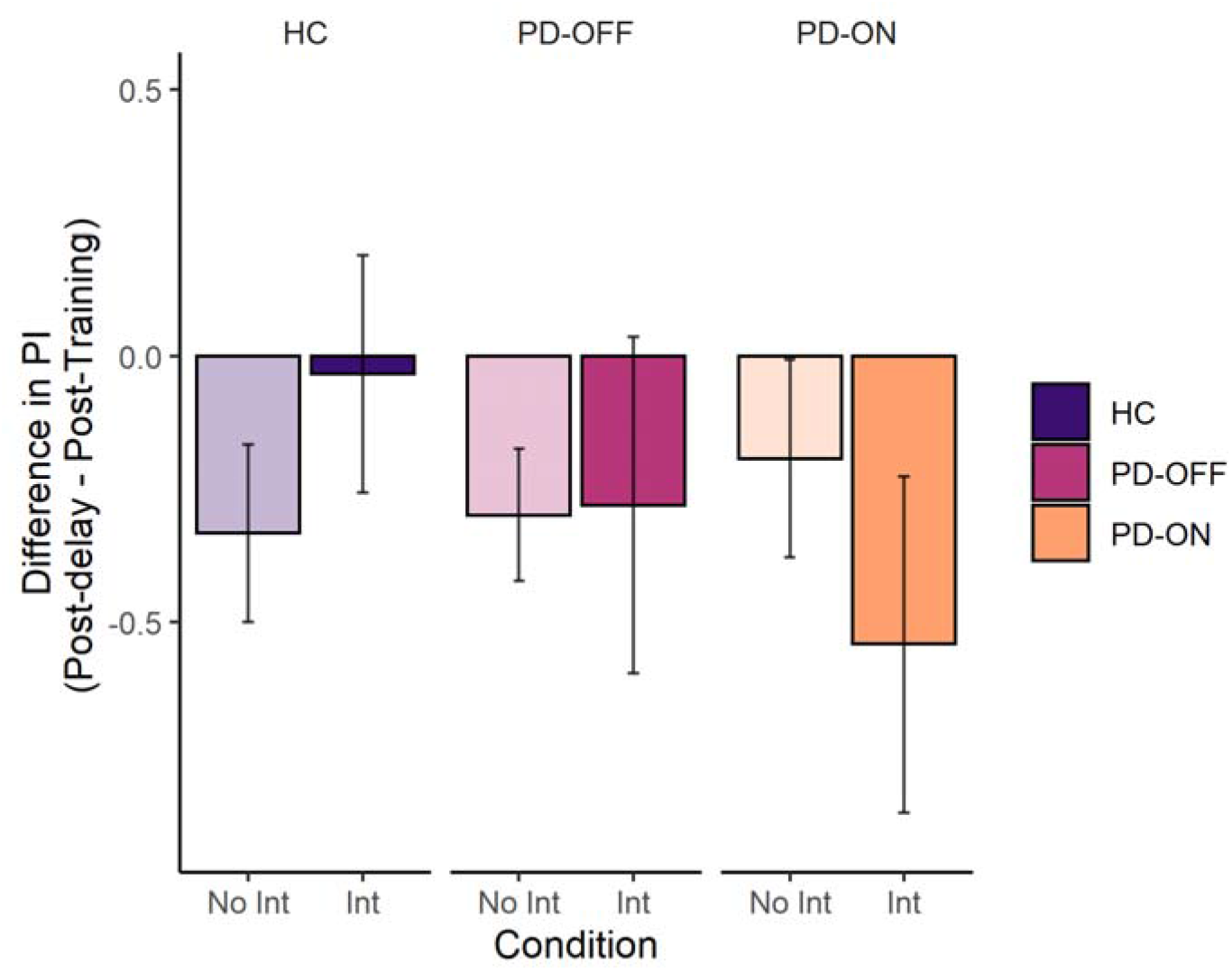
Change in performance across the two-day delay and effect of interference. Change in performance was calculated as the difference between the mean performance at the Post-Delay test and at the Post-Training test. Negative values represent a decay in performance across the delay. There were no differences in the degree of maintenance of performance between groups nor was there an effect of interference on the maintenance of performance. Error bars represent s.e.m. within group and condition. No Int=No interference condition; Int=Interference condition.

A subset of participants was receiving dopamine replacement overnight as part of their regular treatment regimen. Though we did not experimentally manipulate overnight dopamine, the presence of dopamine during sleep could also influence memory. We therefore conducted exploratory analyses to isolate the effect on memory of dopamine during acquisition from the effect of dopamine overnight. First, we focused on the subset of participants from each group who were not receiving any dopaminergic medications overnight. This allowed us to isolate the effect of dopaminergic replacement at the time of acquisition on consolidation. Though this is a smaller sample (14/25 PD-OFF and 11/23 PD-ON), the pattern of results is similar to those obtained in the full sample: performance maintenance across the delay was similar between groups (Group [OFF, ON] x Test [Post-Training, Post-Delay]: F(1, 21)=2.72, p=0.11). Second, to examine the effect of overnight dopamine within each group, we compared patients who were receiving overnight dopamine replacement to patients who were not. In patients ON at the time of initial acquisition, overnight dopamine replacement seemed to benefit performance after the delay (Overnight dopamine [On, Off] x Test [Post-Training, Post-Delay]: F(1, 21)=4.19, p=0.053), but this was not the case in patients OFF (Overnight dopamine [On, Off] x Test [Post-Training, Post-Delay]: F(1,23)=0.50, p=0.48).

### 3.3. Effects of aging on motor memory consolidation

Though the absence of off-line gains in older adults has previously been shown in the literature, the effects of interference on motor memory in older adults have been largely under-explored. We therefore directly compared the different measures of memory between younger and older adults. First, as expected, older adults differed from young adults in the change in performance that occurred across the two-day delay (Group [YC, HC] x Test [Post-Training, Post-Delay]: F(1, 53)=15.20, p=0.0002). The older adults did not show offline gains, but rather maintained their performance at a relatively stable level, as reported above. Furthermore, older adults were less susceptible to the effects of interference on memory than young adults (Group [YC, HC] x Interference [NoInt, Int] x Test [Post-Training, Post-Delay]: F(1, 53)=4.68, p =0.03). As reported above, maintenance of performance across the delay was similar in the older adults who underwent interference and those who did not, whereas younger adults who underwent interference tended to have less offline gains in performance that those who did not.

## 4. DISCUSSION

Motor learning consolidation is known to depend on the striatum and, in keeping with dopamine’s role in supporting corticostriatal plasticity, has generally been thought to depend on dopaminergic inputs to the striatum (for a recent review see Doyon et al., 2018). There is also accumulating evidence from the field of declarative memory or hippocampal-dependent learning, that dopamine state around the time of initial learning plays a role in the later memory for what was learned (Bethus et al., 2010; Chowdhury et al., 2012; Sharp et al., 2020). However, whether dopamine at the time of initial learning of a *motor* skill similarly modulates the later memory for that skill, and whether this process is altered in Parkinson’s patients remains unknown. Here, we investigated whether Parkinson’s patients have impaired memory of motor learning, and whether memory can be facilitated by dopamine replacement at the time of learning. Contrary to our predictions, we found that the degree of motor memory impairment was not greater in patients than in older controls, and was not influenced by dopamine state at the time of initial learning. Specifically, performance was maintained across a two-day delay in patients to the same extent as it was in older controls, yet, in contrast to younger controls, neither the patients nor the older controls showed offline gains in performance. To examine the effects of dopamine and disease on post-learning consolidation processes that are thought to be occurring in the early hours after initial learning, we also used an interference manipulation that has been previously used to probe consolidation processes (Korman et al., 2007). Interestingly, though young controls showed the expected susceptibility to interference delivered 2 hours after learning, patients On or Off dopaminergic medications and older adults did not show a reduction of their memory following interference. These findings suggest that motor memory in Parkinson’s disease, which was preserved in patients to the same degree as in older adults, is supported by compensatory non-dopamine sensitive and possibly extra-striatal mechanisms. Furthermore, given the absence of susceptibility to interference in both patients and older adults, these findings raise the possibility of a shared, age-related mechanism underlying the inability to improve offline.

Consistent with previous work we found that though Parkinson’s patients did not exhibit offline gains, they did maintain their motor memory to the same extent as older adults. Our findings extend this previous work by additionally showing that dopamine state at the time of initial acquisition of the motor skill did not influence memory maintenance, nor the susceptibility to interference. These results are surprising considering the well-established link between motor memory and striatal activity (Albouy et al., 2013b, 2008; Debas et al., 2014, 2010; King et al., 2017), and suggests that Parkinson’s patients are, at least to some extent, relying on non-striatal and non-dopamine-sensitive compensatory mechanisms for the maintenance of motor memories. Recruitment of compensatory processes in Parkinson’s patients has been well described (Appel-Cresswell et al., 2010; Palmer et al., 2010; Yu et al., 2007). One possible compensatory substrate is the cerebellum, which has been proposed as a region that plays a compensatory role for both motor and non-motor processes in Parkinson’s disease (Wu & Hallett, 2013). Several functional neuroimaging studies in Parkinson’s patients using a variety of motor tasks have demonstrated increased task-related activity in the cerebellum (Palmer et al., 2009; Wu and Hallett, 2005; Yu et al., 2007) as well as increased functional connectivity of the cerebellum in the setting of relatively normal motor performance (Festini et al., 2015; Mentis et al., 2003; Palmer et al., 2009; Simioni et al., 2016; Wu et al., 2009). For instance, in Parkinson’s patients OFF medication, better motor performance was associated with increased motor-task-related BOLD activity in the cerebellum (Palmer et al., 2009) and increased cerebellar-putamen functional connectivity (Simioni et al., 2016). No study has specifically investigated the role of the cerebellum in the process of motor memory in Parkinson’s patients. However, given evidence that it is recruited to a greater degree during execution of simple motor tasks in Parkinson’s patients, it is plausible that the cerebellum may also be involved during the initial acquisition of a motor skill, which may render the subsequent process of memory independent of the striatum and, in the case of Parkinson’s disease patients, independent of their dopamine state. Another possible substrate for compensatory motor memory is the hippocampus. The hippocampus remains relatively spared in the earlier stages of Parkinson’s disease and in patients who are free of dementia, as was the case for the patients included in our sample (Hawkes et al., 2010) and could therefore potentially be recruited to support motor memory. Indeed, several studies have shown that the striatum and the hippocampus both support the consolidation of motor sequence memories (Albouy et al., 2013b, 2008), though it has been proposed that they support distinct aspects of consolidation on this task (Albouy et al., 2015).

We did not find evidence that dopamine at the time of the initial acquisition of motor learning influences the subsequent memory of that learning. Our motivation for choosing to focus on dopamine at the time of acquisition was informed by evidence that the degree of involvement of the striatum at the time of initial learning predicts offline consolidation and later memory (Albouy et al., 2013b, 2008; King et al., 2017). This is also consistent with models of corticostriatal plasticity, which propose that dopamine release in the period surrounding activation of a synapse influences plasticity (Calabresi et al., 2007; Wickens, 2009). However, evidence from the study of episodic, or hippocampal-dependent memory, which also shows a clear beneficial effect of dopamine at the time of initial encoding on later memory, specifically shows that this effect is tied to the interaction between dopamine and an environmental signal such as reward (McNamara et al., 2014; Redondo and Morris, 2011; Sharp et al., 2020; Shohamy and Adcock, 2010; Wang et al., 2010). We did not manipulate reward during learning. Thus, it is possible that simply restoring dopamine is not sufficient, and that an environmental trigger for dopamine release is also necessary for a beneficial effect on memory to occur. Indeed, several recent studies of motor learning, including one that relied on a sequence learning task similar to ours (Wächter et al., 2009), have shown that reward can enhance the retention of motor memories in healthy controls (Abe et al., 2011; Galea et al., 2015). Future work will be necessary to determine whether the combination of reward and dopamine replacement could enhance motor memory in Parkinson’s patients, or whether reliance on extra-striatal compensatory mechanisms eliminates these potential beneficial effects.

Dopamine state during the early post-encoding period and during overnight sleep may also be important but very little research exists to guide hypotheses. Given the 1.5-2-hour half-life of levodopa, it is reasonable to assume that patients ON were still ON in the early post-encoding period, whereas the patients OFF remained OFF. Given that patients ON and OFF did not differ in their memory maintenance, this suggests that dopamine state during the early post-encoding period is not a key modulator of subsequent consolidation. Exploratory analyses examining the effect of overnight dopamine within each group, comparing patients who were receiving dopaminergic medications overnight to those who were not revealed a possible benefit of overnight dopamine state on consolidation but only in patients who were also ON at the time of initial acquisition, which raises the possibility of an interaction between the neural processes underlying the initial learning and those underlying the later consolidation process. Future studies specifically manipulating dopamine at the different critical periods of the consolidation process are required.

An unexpected finding in our study is the fact that memory maintenance in older adults was not susceptible to interference. The older adults therefore differed from the younger adults in two key ways: they did not show offline gains, and did not show susceptibility to interference. Previous work has similarly demonstrated the absence of offline gains in older adults (Fogel et al., 2014; Spencer et al., 2007; Wilson et al., 2012), and has suggested that this age-related impairment in consolidation is due to age-related changes in sleep, such as reduced spindles (Fogel et al., 2014; Harand et al., 2012; King et al., 2013; Wilson et al., 2012). However, the absence of a susceptibility to interference in older adults points to an additional mechanism. Specifically, our results suggest that the consolidation processes that usually take place in the first few hours following learning - while the memory is still labile, and which are thought to be blocked by interference (Korman et al., 2007; Walker et al., 2003), are either compromised in older adults and patients (Korman et al., 2015), or are occurring at a different, possibly later, timepoint. Whether the mechanism underlying this change is similar in the patients and the controls remains to be determined, but recent evidence showing that corticostriatal networks are indeed important even at this early post-learning stage (Censor et al., 2014), and are altered in older adults across the consolidation period (Fogel et al., 2014), suggests that the cause of disrupted consolidation may be shared.

In conclusion, our findings demonstrate that motor memory in Parkinson’s patients is similar to that of older adults in that patients were able to maintain their performance but did not show offline gains. Furthermore, we showed that neither patients nor older adults were susceptible to an interference manipulation. These similarities suggest that changes in motor memory seen in Parkinson’s disease might be explained by age-related mechanisms (King et al., 2013; Korman et al., 2015), suggesting a shared neural substrate for motor memory across groups. We also found that dopamine at the time of initial learning did not influence later memory for that learning. This suggests a reliance on extra-striatal and non-dopamine-sensitive networks for motor memory in Parkinson’s patients and, given the similar pattern of performance observed in the patients and older adults, could provide clues for identifying the mechanisms that underlie age-related changes to motor memory. An important future step will be to leverage neuroimaging to identify key compensatory mechanisms and begin to understand the evolution of compensation over the course of aging and disease, as well as factors that influence the success of such compensation. Future work will also be required to establish, in both healthy aging and Parkinson’s disease, the mechanisms that affect memory consolidation across the full timeline of this process, from the point of initial learning to delayed retrieval. We found an absence of susceptibility to early interference in both the older adults and the patients, suggesting that the process of memory transformation is already altered in the first few hours following learning. Sleep alterations, believed to contribute to motor consolidation deficits in aging, may represent an even more important mechanism in patients (Latreille et al., 2016, 2015). Identifying such factors will be important as there is evidence for effective sleep therapies in Parkinson’s patients (Gros et al., 2016; Kaminska et al., 2018; McCarter et al., 2013) and even recent evidence that dopamine state may impact the relationship between sleep and memory (Feld et al., 2014; Isotalus et al., 2020).

## Acknowledgements

We thank Rebecca Kahane for help with participant testing, Drs. Roy Alcalay and Cheryl Waters for help with patient recruitment, and the participants for their time and interest. This work was supported by a pilot award from the Parkinson’s Foundation. MS was supported by Healthy brains, Healthy Lives. SL was supported by the Fonds de Recherche du Québec – Santé.

