## Supplementary_material for "Preserved motor memory in Parkinson’s disease"

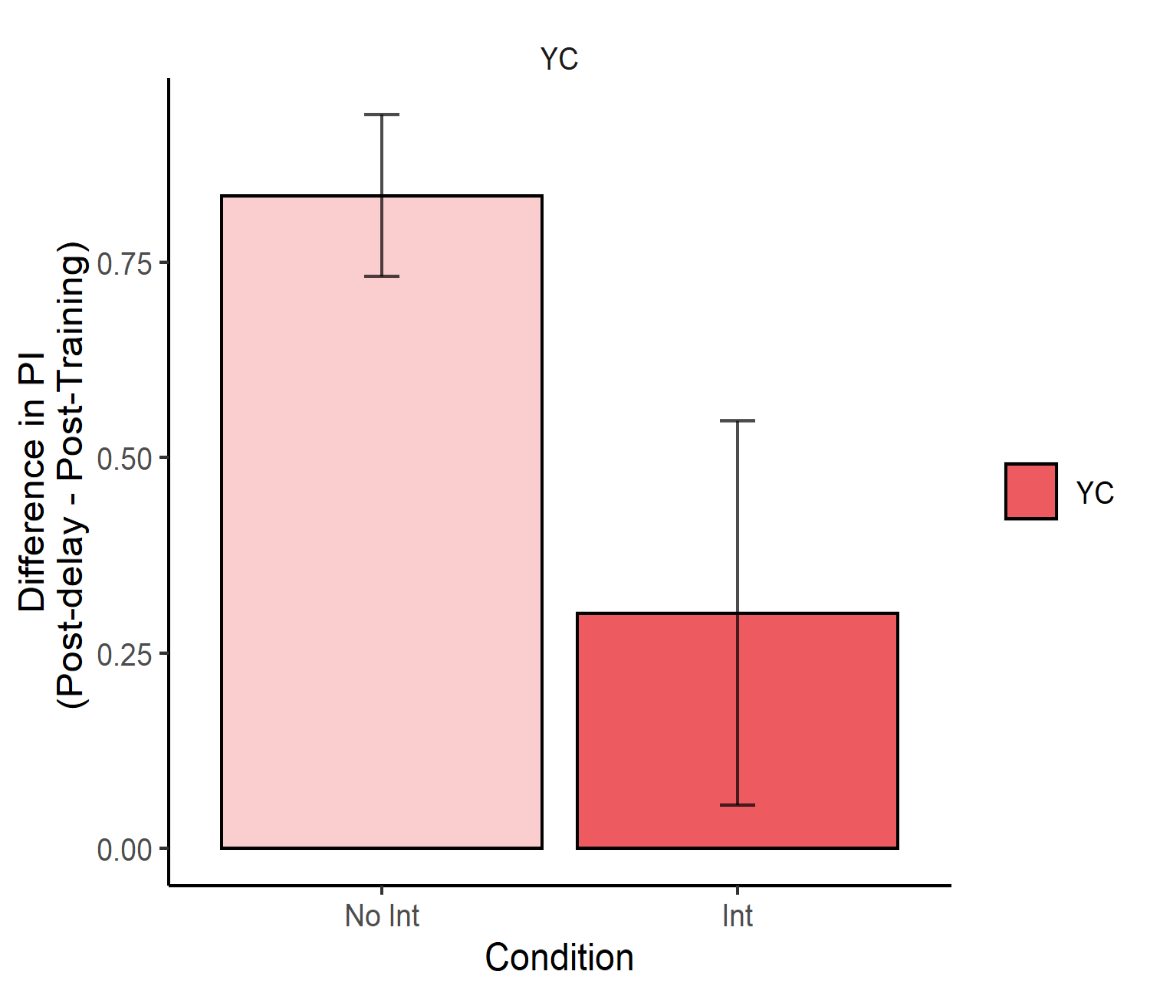

**Supplemental figure 1. Change in performance across the two-day delay and effect of interference among the young controls.** Change in performance was calculated as the difference between the mean performance index at the Post-Delay test and at the Post-Training test. Negative values represent a decay in performance across the delay and positive values represent gains in performance. Young controls showed an off-line improvement in performance across the delay, and this improvement was significantly reduced in the group that underwent interference (*: t = 2.00, p = 0.05). Bars represent s.e.m. within group and condition. No Int=No interference condition; Int=Interference condition.

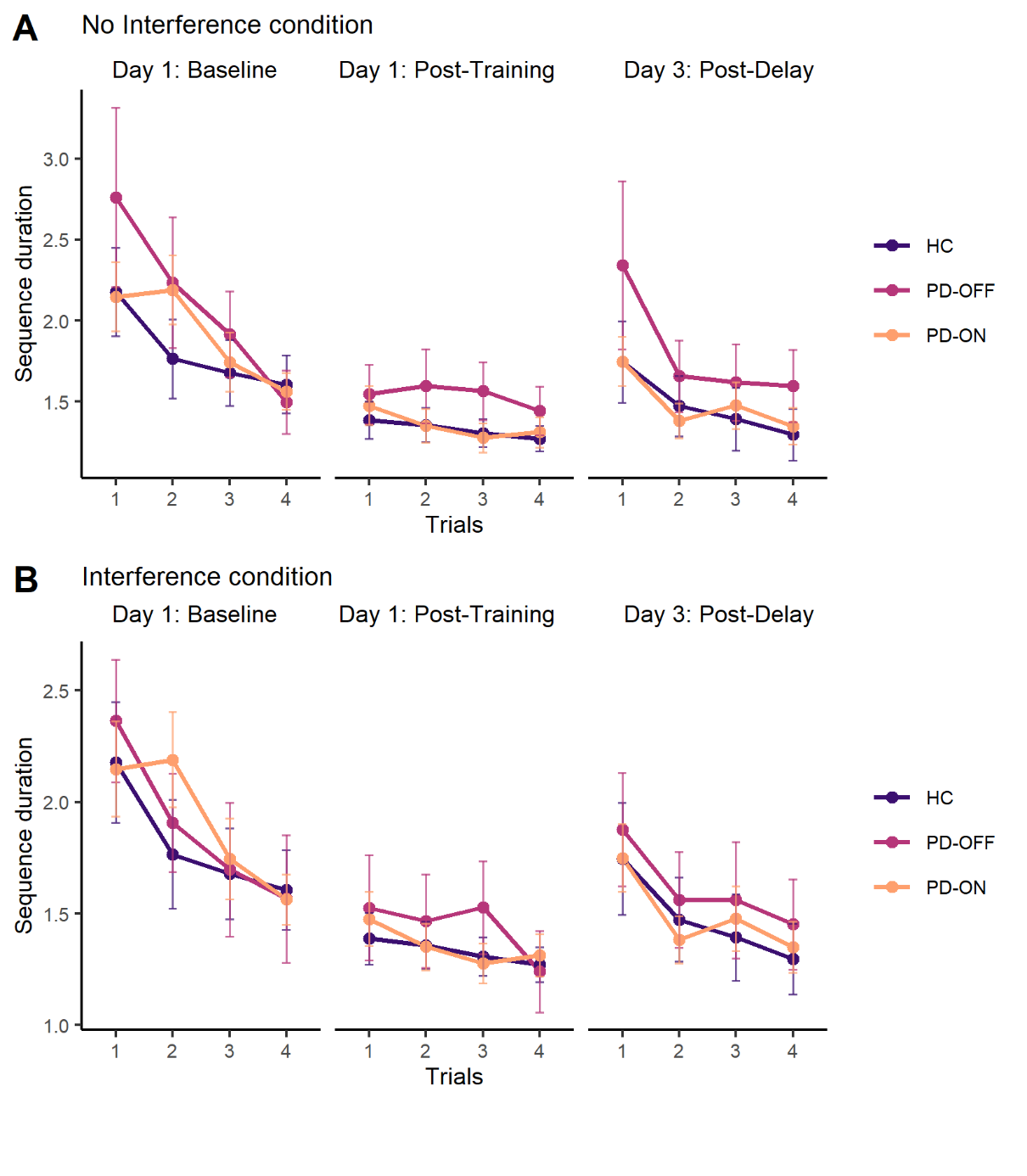

**Supplemental figure 2. Test performance (Sequence duration).** Performance measured as the average sequence duration is shown across trials (1-4) for each of the three test sessions for all three groups. Conditions are presented separately: participants that did not undergo the interference (A) and those that did (B). Participants across groups and conditions show similar baseline performance, similar improvements from training and similar decay in performance across the two-day delay. Error bars represent s.e.m. within group and trial.

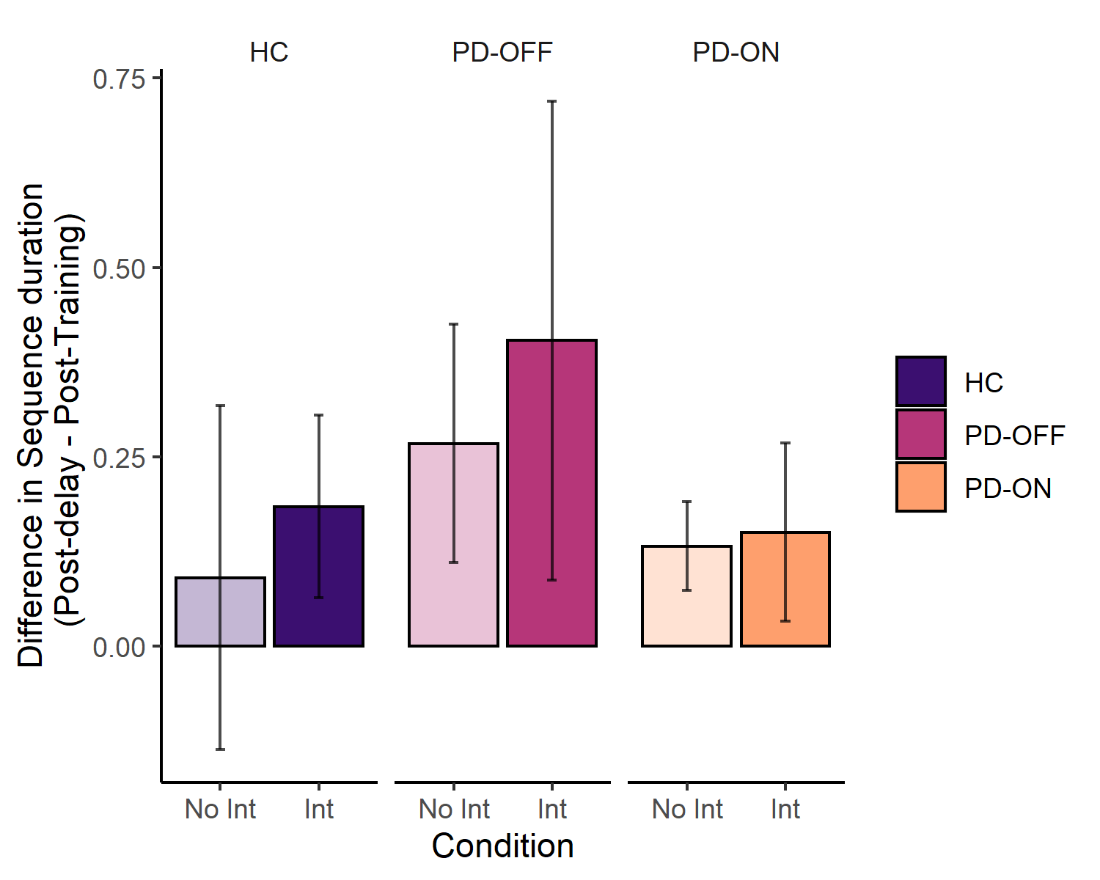

**Supplemental figure 3. Change in performance across the two-day delay and effect of Interference on mean sequence duration.** Change in performance was calculated as the difference between mean sequence duration at the Post-Delay test and at the Post-Training test. Error bars represent s.e.m. within group and condition Negative values represent a decay in performance across the delay. There were no differences in the degree of maintenance of performance between groups nor was there an effect of interference on the maintenance of performance. No Int=No interference condition; Int=Interference condition.

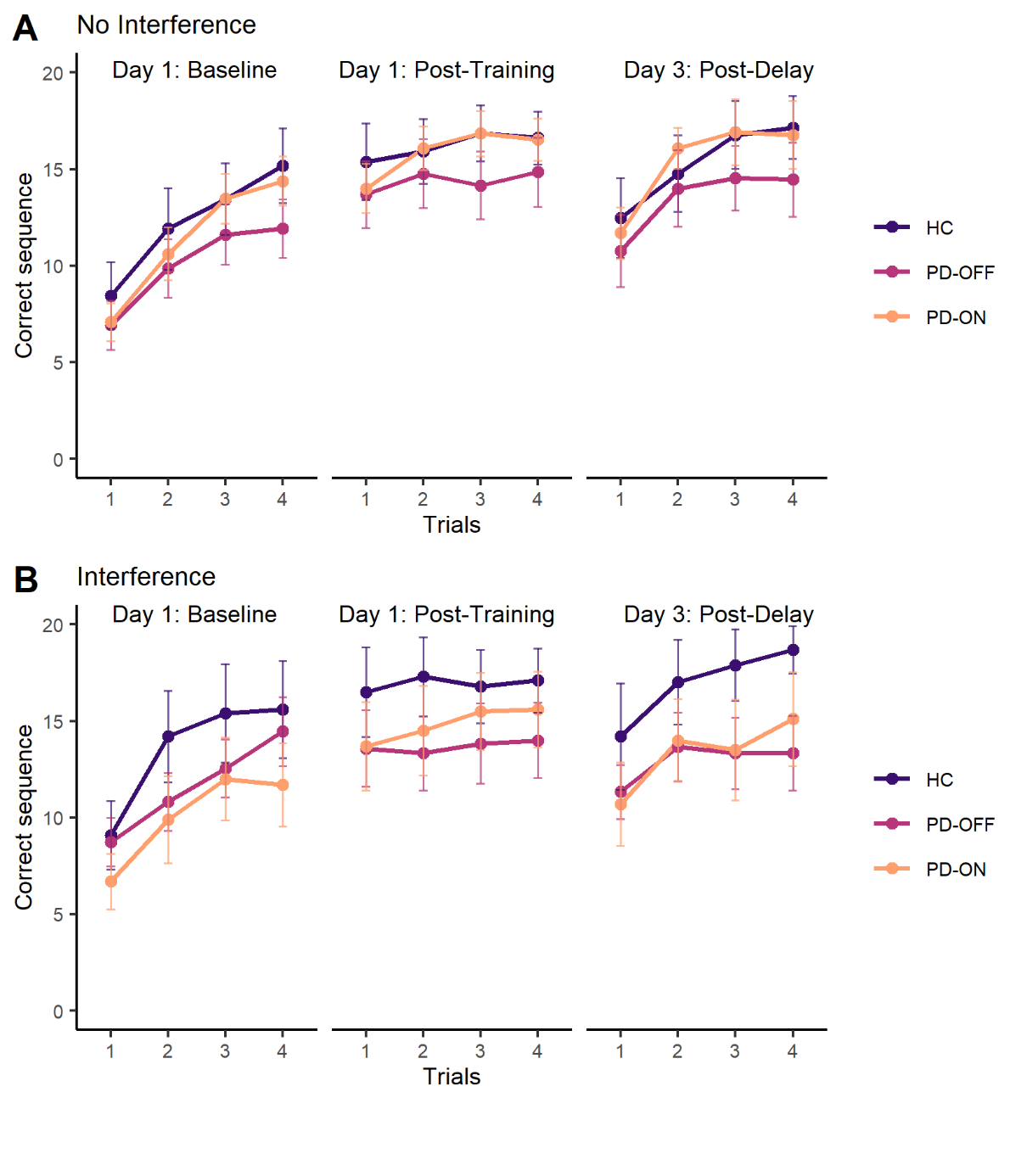

**Supplemental figure 4. Test performance (Correct sequences).** Performance measured as the average number of correct sequences is shown across trials (1-4) for each of the three test sessions for all three groups. Conditions are presented separately: participants that did not undergo the interference (A) and those that did (B). Error bars represent s.e.m. within group and trial.

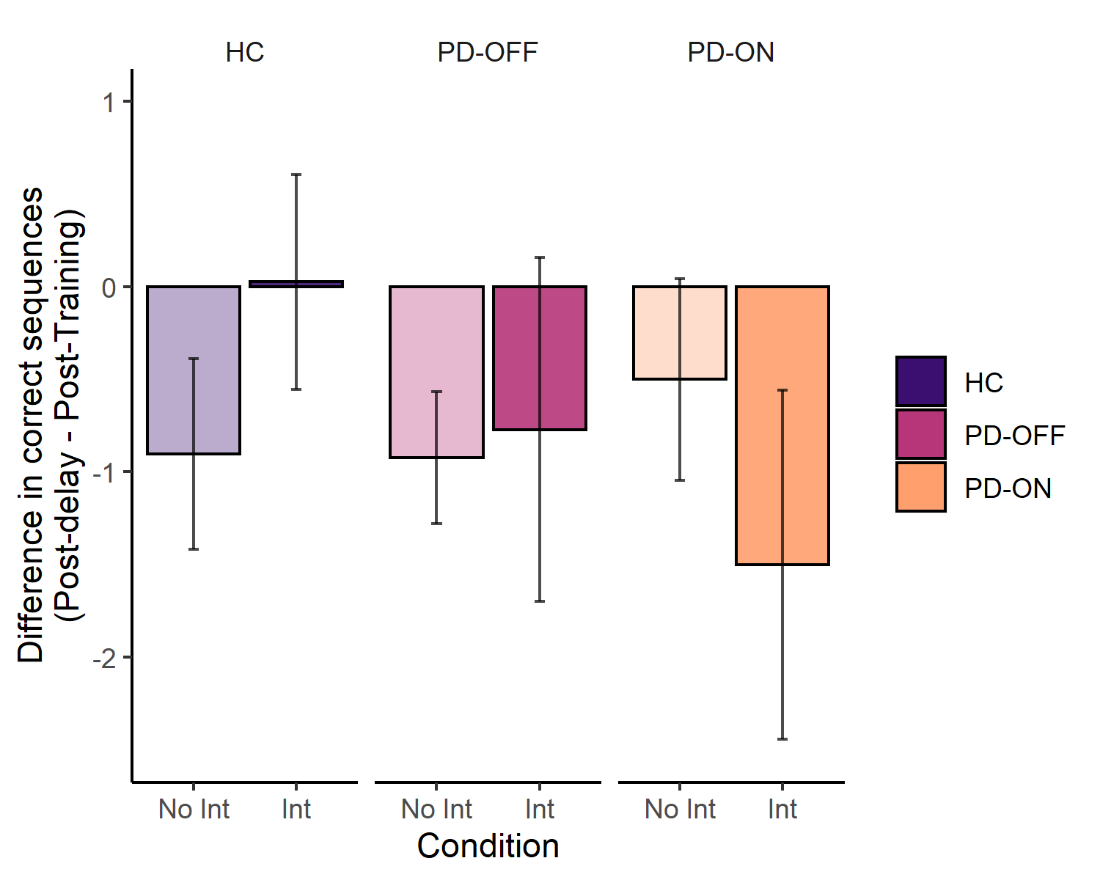

**Supplemental figure 5. Change in number of correct sequences across the two-day delay and effect of Interference.** Change in performance was calculated as the difference between average number of correct sequences at the Post-Delay test and at the Post-Training test. Error bars represent s.e.m. within group and condition. No Int=No interference condition; Int=Interference condition.
